# Identification of a peroxidase inhibitor that enhances kanamycin activity

**DOI:** 10.1101/2020.10.12.336933

**Authors:** Zhen Hui, Shiyi Liu, Ruiqin Cui, Biao Zhou, Chunxia Hu, Min Zhang, Qiuyang Deng, Shumin Cheng, Yutian Luo, Huaisheng Chen, Jinsong Wu, Yuemei Lu, Xueyan Liu, Lingyun Dai, Wei Huang

## Abstract

**Background:** The threat of antimicrobial resistance calls for more efforts in basic science, drug discovery, and clinical development, particularly gram-negative carbapenem-resistant pathogens.

**Objectives and methods:** Whole-cell–based screening was performed to identify novel antibacterial agents against *Acinetobacter baumannii* ATCC19606. Spontaneously resistant mutant selection, whole-genome sequencing, and surface plasmon resonance were used for target identification and confirmation. Checkerboard titration assay was used for drug combination analysis.

**Results:** A small molecule named 6D1 with the chemical structure of 6-fluorobenzo[d]isothiazol-3(2H)-one was identified and exhibited activity against *A. baumannii* ATCC19606 strain (minimal inhibitory concentration, MIC = 1 mg/L). The mutation in the plasmid-derived *ohrB* gene that encodes a peroxidase was identified in spontaneously resistant mutants. Treatment of the bacteria with 6D1 resulted in increased sensitivity to peroxide such as *tert*-butyl hydroperoxide. The binding of 6D1 and OhrB was confirmed by surface plasmon resonance. Interestingly, the MIC of kanamycin against spontaneously resistant mutants decreased. Finally, we identified the effect of 6D1 on enhancing the antibacterial activity of kanamycin, including New Delhi metallo-β-lactamase (NDM-1)-producing carbapenem-resistant *Klebsiella pneumoniae*, but not in strains carrying kanamycin resistance genes.

**Conclusions:** In this study, we identified a peroxidase inhibitor that suppresses the growth of *A. baumannii* and enhances the antibacterial activity of kanamycin. We propose that peroxidase may be potentially used as a target for kanamycin adjuvant development.

## Introduction

Gram-negative pathogens such as *Acinetobacter baumannii*, *Klebsiella pneumoniae*, and *Pseudomonas aeruginosa* have become resistant to almost all commonly used antimicrobial agents, including aminoglycosides, quinolones, and broad-spectrum β-lactams. Overall, for instance, approximately 45% of all global *A. baumannii* isolates are considered multidrug-resistant (MDR) (bacteria resistant to more than three antibiotic classes).^1,2^

With the emergence of carbapenem-resistant gram-negative pathogens such as *A. baumannii* (CRAB), tigecycline and polymyxin-class antibiotics are the only currently available treatment options.^3^ However, treatment outcomes of tigecycline have been hampered by the low serum concentrations of the drug in the approved dosing regimen and the low penetration in the epithelial lining fluid (ELF) of mechanically ventilated patients.^4^ The resistance of polymyxin-class antibiotics as well as nephrotoxicity and neurotoxicity are the major factors that limit the usage of polymyxin. Previous clinical observations showed that the rates of nephrotoxicity occurred in approximately 60% of patients who received colistin or polymyxin B therapy.^5–7^ Eravacycline, cefiderocol, and plazomicin seem to be promising new agents against *A. baumannii*. However, evaluation of their position in clinical practice and particularly in ventilator-associated pneumonia (VAP) has not been performed to date.^8,9^

The present clinical pipeline does not meet current needs, and thus more investment is required in basic science, drug discovery, and clinical development, particularly gram-negative carbapenem-resistant pathogens, including CRAB.^10^ Therefore, we launched a whole cell-based screening program for *A. baumannii*. Here, we report the discovery of compound 6D1 that exhibits anti-*A. baumannii* activity. In addition, we show that enhancement effect of 6D1 on the antibacterial efficacy of kanamycin through the inhibition of plasmid-derived OhrB.

## Materials and methods

### Bacterial strains, growth conditions, reagents, and screening strategy

*A. baumannii*, *K. pneumoniae*, and *P. aeruginosa* were grown in liquid broth (LB) medium or LB agar. Antibiotic (purchased from Sigma-Aldrich, USA) solutions were prepared at a concentration of 1 mg/mL in distilled water or 100% dimethylsulfoxide (DMSO), filter-sterilized, and frozen at −20°C until use. The compounds to be screened were dissolved in 100% DMSO and stored as frozen stocks at a concentration of 1 mg/mL.

We sought anti-*A. baumannii* compounds by testing compounds for inhibition of *A. baumannii* ATCC19606. A whole-cell assay was used because of its ability to concurrently assess multiple targets. Compounds were prepared in 96-well plates at a concentration of 10 mg/L in 50 μL LB broth. A 50-μl aliquot of each bacteria culture was then added to each well of the 96-well plate at an OD_600_ = 0.006. The plates were incubated overnight at 37^°^C, and the primary active hits were filtered by achieving at least 90% of bacterial growth inhibition using Cell Counting Kit-8 (MCE, USA). Subsequently, two-fold serial dilutions of primary hits were prepared for the determination of minimal inhibitory concentration (MIC, defined as the lowest concentration of compound that inhibited 90% of bacterial growth). Compounds with an MIC ≤ 1 mg/L were selected for further investigation.

### Spontaneously resistant mutant selection

Spontaneously resistant mutants were selected via stepwise exposure to increasing concentrations of the compounds. An aliquot of mid-log phase (OD_600_ = 0.6) bacterial culture (1 mL) was added to 2 mL of medium containing serial increasing concentrations of 6D1 until no growth was observed. The bacteria that survived in culture were spread onto agar plates containing the corresponding concentrations of the 6D1 compound. All colonies that originated from different plates and represent independent biological events were subjected to whole-genome sequencing (WGS). The resistance phenotype to the compound was confirmed by testing for a shift in MIC values.

### WGS

Genomic DNA was extracted from each isolate using a gram-negative bacterial genome extraction kit (Tiangen, China). Whole-genome fragment libraries were prepared using a paired-end sample preparation kit (Illumina, USA). The genomes were sequenced using Illumina HiSeq 2500 platform (Illumina, USA) and assembled with *de novo* SPAdes Genome Assembler (version 3.12.0).^11^ The resulting reads were mapped to the *A. baumannii* ATCC19606 reference genome, and mutations were identified using Snippy (https://github.com/tseemann/snippy).

### Effect of compounds on the tolerance of bacteria to peroxides

The effects on the tolerance of bacteria to peroxides was determined by testing for shifts in MIC of *tert*-butyl hydroperoxide (*t*-BHP), cumene hydroperoxide (CHP), and hydrogen peroxide (H_2_O_2_) in the presence of the compound.

### Protein expression and purification

The cDNA encoding for full-length OhrB was chemically synthesized with codon optimization for expression in *E. coli*. Vector pET28b was used for protein expression. The plasmid pET28b-*ohrB* was then transformed into competent BL21 strain cells. The BL21 cells carrying the aforementioned plasmid were grown in LB medium at 37^°^C to an OD_600_ of 0.4. The cell cultures were then supplemented with 0.5 mM isopropyl β-D-1-thiogalactopyranoside (IPTG). The induced cells were further grown at 16^°^C with shaking at 220 rpm overnight to induce the expression of the recombinant protein.

To purify the recombinant proteins, the cells were harvested and lysed by ultrasonication. The supernatant of the lysed cells was loaded onto Ni-NTA columns (Qiagen, Germany), and the proteins were further purified by a gel-filtration column (GE Healthcare, USA) with a gel-filtration buffer [100 mM NaCl, 10 mM Tris-HCl (pH 7.5) and 1 mM DTT]. The protein concentration was determined using the Bradford method.

### Surface plasmon resonance (SPR) experiment

OhrB were covalently immobilized to a sensor chip CM5 (29-1049-88, Sweden) by means of amino coupling. The running buffer used in the experiment contains 20 mM HEPES pH 7.5, 150 mM NaCl, 0.05% Tween 20, 0.1% DMSO, and the 6D1 compound was also dissolved in the running buffer. The sensor chip was washed with running buffer between each concentration. Reference runs were performed with blank (sensor chip only) and active (sensor chip with OhrB only) channel on the same sensor chip. The assay curves were constructed using serial concentrations of 6D1 of 7.5, 15, 30, 60, and 120 μM. The kinetic parameters of the interaction and the affinity constants were calculated using Biacore T200 evaluation software.

### Checkerboard titration assay

Drug interactions between 6D1 and the bactericidal drugs were performed using a chequerboard titration assay.^12^ The fractional inhibitory concentration (FIC) was calculated using the following formula: (MIC of drug A or B in combination)/(MIC of drug A or B alone). The fractional inhibitory concentration index (FICI) was determined by adding the two FICs. Synergy, antagonism, and no interaction were defined as FICI ≤ 0.5, FICI > 4.0, and FICI = 0.5–4.0, respectively.^12^

## Results

### *In vitro* activity of 6D1

We identified an active hit named 6D1 (MIC = 1 mg/L) with the structure of 6-fluorobenzo[d]isothiazol-3(2H)-one, which is similar to 1, 2‐benzisothiazolin‐3‐one (BIT) and an antifungal drug ticlatone (Figure 1). A moderate antibacterial activity of 6D1 was also observed in *S. aureus* (MIC = 2.5 mg/L), but not in *K. pneumoniae*, and *P. aeruginosa* (MIC ≥ 5 mg/L). Unexpectedly, MICs of 6D1 were high in CRAB clinical isolates (MIC = 5–10 mg/L) (Table 1).

**Table 1.**
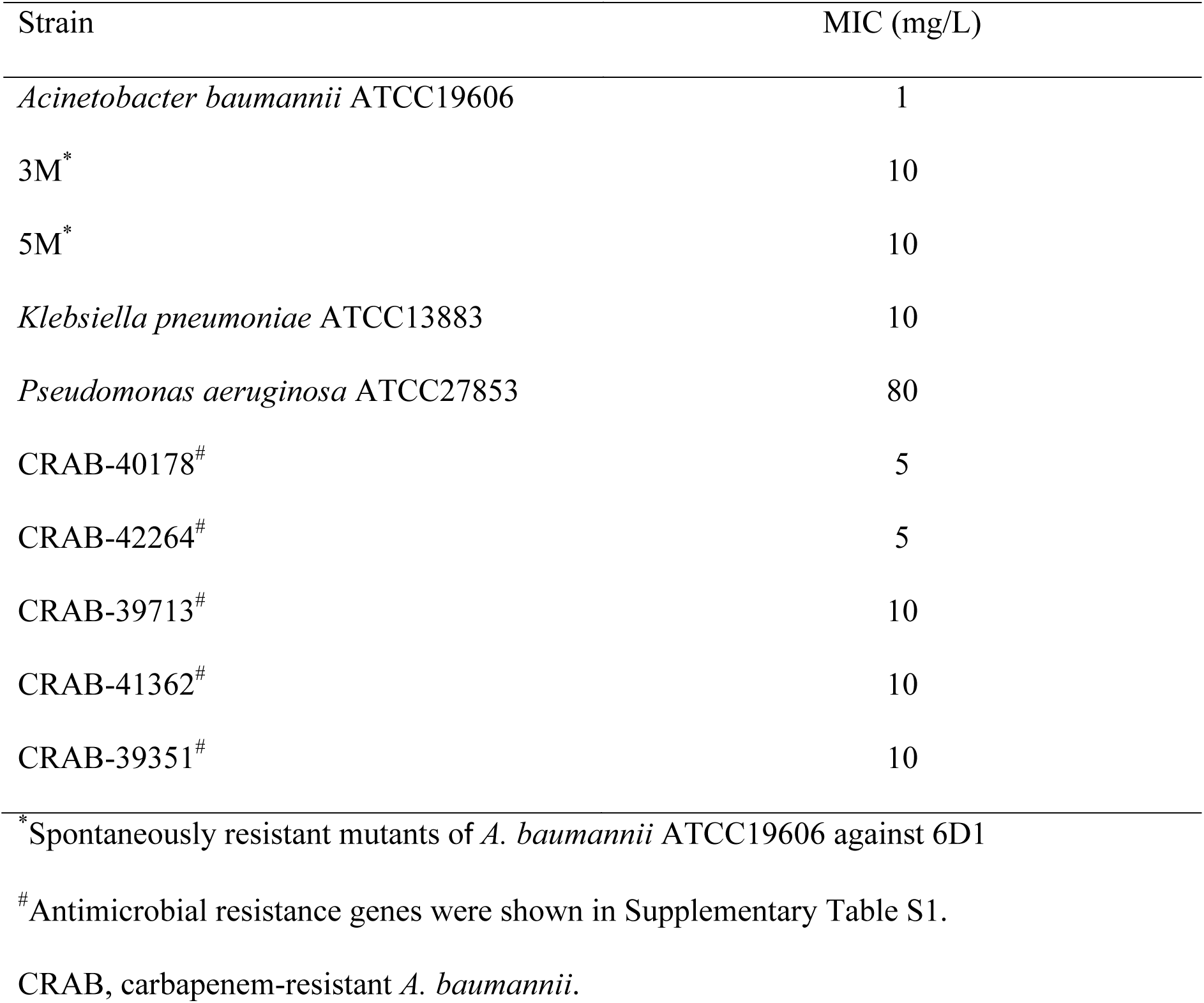
MICs of 6D1 that inhibited 90% of the growth of different bacterial strains

**Figure 1.**
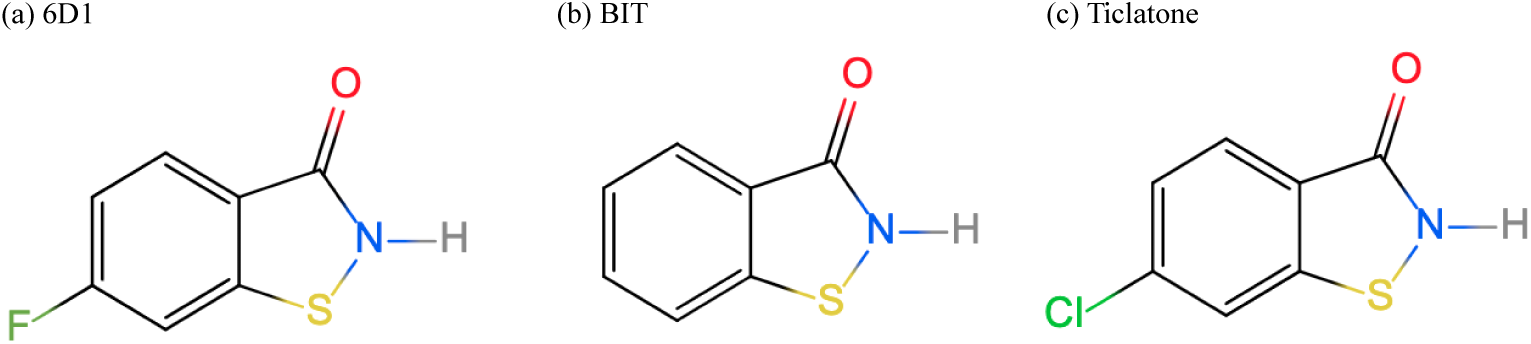
Chemical structure of 6D1 and its analogues. (a) 6D1. (b) 1, 2‐benzisothiazolin‐3‐ one (BIT). (c) Ticlatone.

### OhrB mutations confer resistance to 6D1

To identify the target of 6D1, we obtained two 6D1 spontaneously resistant strains (3M and 5M) from independent cultures with bacterial growth in LB broth containing 10× MIC (10 mg/L) of 6D1. An increase in MIC indicated the resistance phenotype of 3M and 5M to 6D1 (Table 1). Compared with the wild-type (WT) parent strain *A. baumannii* ATCC19606, mutations located in plasmid (pMAC)-derived *ohrB* were identified both in 3M and 5M strains, resulting in the conversion of arginine at the position 15 (Arg15) of OhrB (Table 2). Molecular dynamics simulations and *in silico* mutagenesis indicated that the corresponding Arg19 in Ohr from *Xylella fastidiosa* contributed to the stabilization of XfOhr in the closed state, suggesting that the mutations in 3M and 5M probably affect the function of OhrB.^13^

**Table 2.** Whole-genome sequencing identified polymorphisms within spontaneously resistant mutants of *A. baumannii* ATCC19606 against 6D1

| Position of A.<br>*baumannii* 19606 | Reference | 3M | 5M | Locus<br>(DJ41_) | Gene | Product | Effect of mutation |
| --- | --- | --- | --- | --- | --- | --- | --- |
| NZ_KL810966.1:<br>1913491 | G | T | - | RS13465 | - | LysR family transcriptional regulator | Asn272Lys |
| NZ_KL810966.1:<br>2417811 | G | - | A | RS0104185 | - | hypothetical protein | Gly951Asp |
| NZ_KL810967.1:<br>13770 | C | - | T | RS22870 | *ohr* | organic hydroperoxide resistance protein | Arg15His |
| NZ_KL810967.1:<br>13771 | G | A | - | RS22870 | *ohr* | organic hydroperoxide resistance protein | Arg15Cys |

### 6D1 reduces the tolerance of bacteria to peroxides

Ohr was first described in *Xanthomonas campestris*. It has since been found in a number of bacterial species.^14, 15^ Owing to the Cys-based, thiol-dependent peroxidase activity, Ohr plays a central role in bacterial responses against fatty acid hydroperoxides and peroxynitrite, thus resulting in an “organic hydroperoxide resistance” phenotype.^16^ Table 3 shows that the MICs of *t*-BHP, CHP, and H2O2 in *A. baumannii* ATCC19606 were at least eight-fold lower when in the presence of 2.5 mg/L of 6D1. The magnitude of MIC reduction coincided with the substrate preference of Ohr (H2O2 <<<< CHP < *t*-BHP) as previously reported.^16^ In contrast, in 3M strain, the MICs of *t*-BHP, CHP, and H_2_O_2_ were almost not affected by the presence of 6D1; whereas in 5M strain, the MICs of *t*-BHP, CHP, and H_2_O_2_ were reduced, indicating that 6D1 could still affect the function of OhrB in 5M strain. However, the MICs of *t*-BHP and H_2_O_2_ in 3M or 5M strains were slightly lower than the WT, suggesting that the *ohrB* mutation resulted in reduced tolerance to peroxides. Moreover, growth retardation was also observed in 3M and 5M strains, but the growth of 3M strain was not affected by 6D1 (Figure 2).

**Table 3.** Effect of 6D1 on the susceptibility to peroxides

| Peroxide<br>(mM) | Wild-type | Wild-type<sup>a</sup> | 3M | 3M<sup>a</sup> | 5M | 5M<sup>a</sup> |
| --- | --- | --- | --- | --- | --- | --- |
| *t*-BHP | 0.2 | <=0.005 | 0.1 | 0.1 | 0.1 | 0.1 |
| CHP | 0.2 | <=0.01 | 0.2 | 0.2 | 0.2 | 0.08 |
| H<sub>2</sub>O<sub>2</sub> | 0.8 | <=0.1 | 0.5 | 0.4 | 0.6 | 0.3 |

<sup>a</sup>The MICs of peroxides in the presence of 2.5 mg/L of 6D1.
*t*-BHP, *tert*-butyl hydroperoxide; CHP, cumene hydroperoxide; H<sub>2</sub>O<sub>2</sub>, hydrogen peroxide.

**Figure 2.**
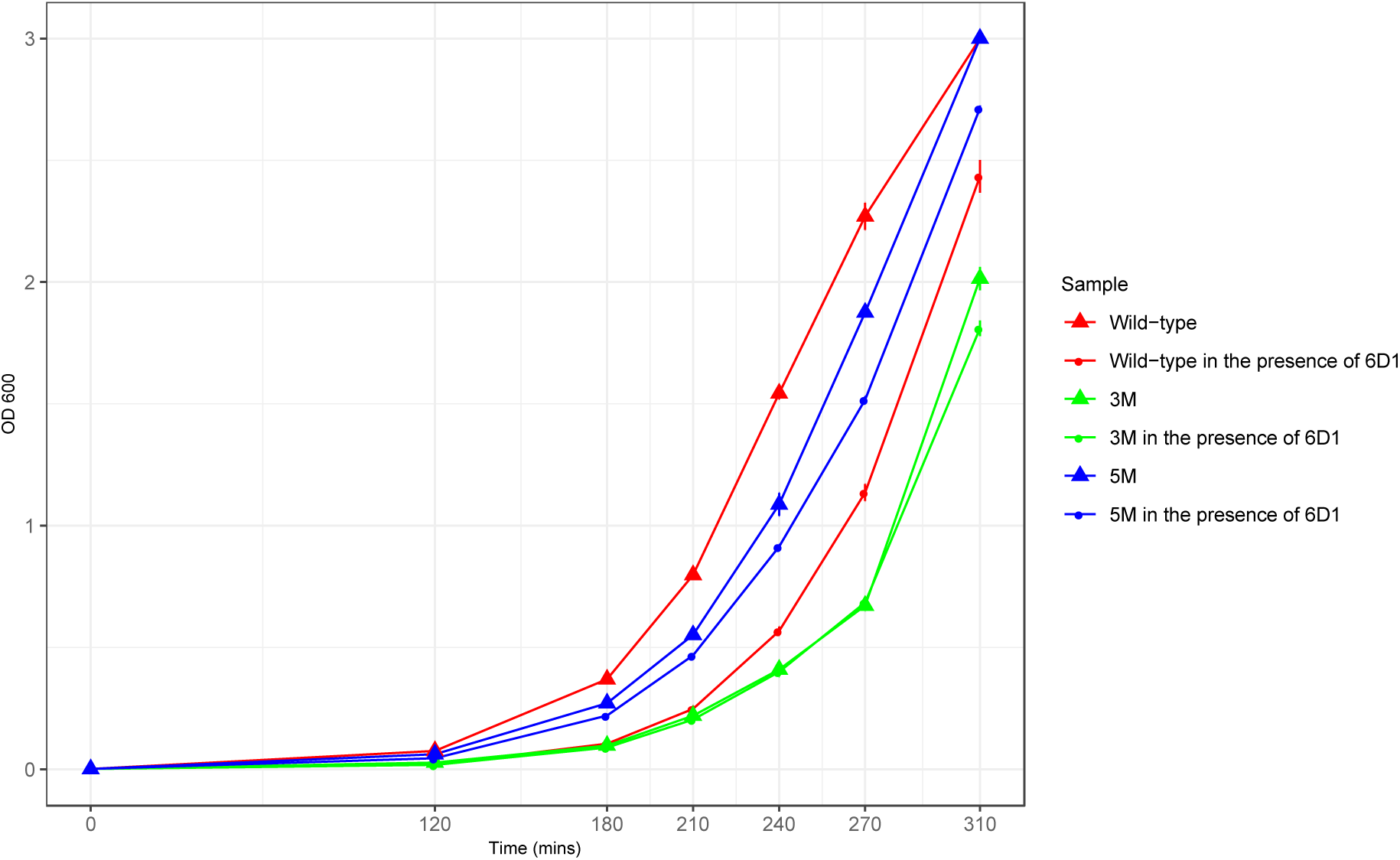
Growth curve of wild-type *Acinetobacter baumannii* ATCC19606, 3M, and 5M in the absence or presence of 0.25 mg/L 6D1.

### MICs of bactericidal drugs in 3M and 5M strains

With aim to identify the effects of OhrB protein function on the activity of bactericidal agents, we tested the MICs of a handful of bactericidal drugs in the 3M, 5M, and WT strains. Compared with the WT, the MICs of kanamycin in 3M and 5M strains decreased by at least 2-fold, thereby suggesting the association between OhrB function and kanamycin activity (Table 4).

**Table 4.**
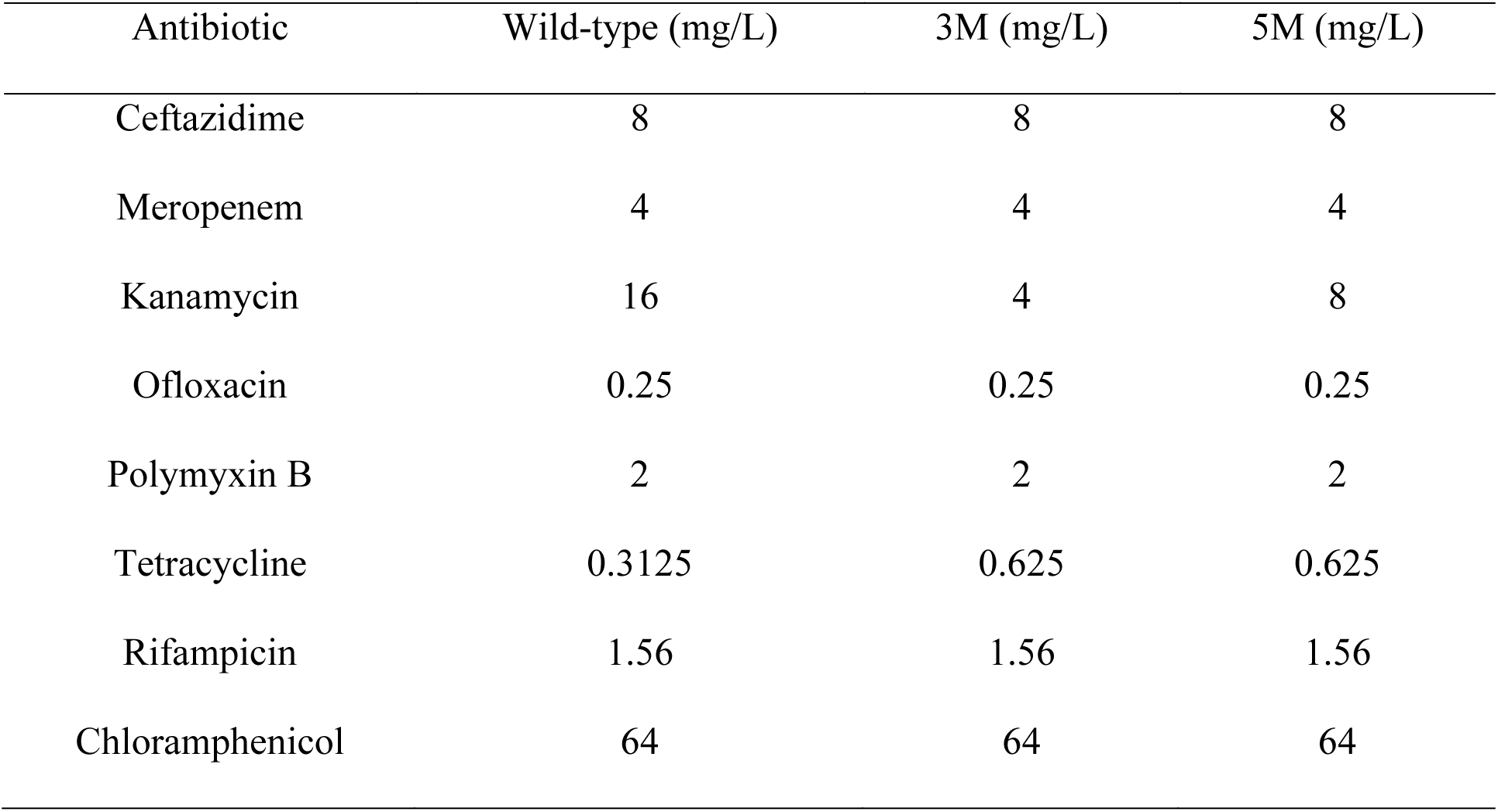
MICs of different classes of antibiotics that inhibited 90% of bacterial growth

### Drug combinations study

The observed changes of MIC for kanamycin in 3M and 5M suggest that 6D1 probably enhances antibacterial activity. Therefore, we used a checkerboard titration assay to identify the drug interaction of 6D1 and kanamycin in different species. Table 5 revealed a synergistic effect between 6D1 and kanamycin in *K. pneumonae* and *A. baumannii* (FICI = 0.5). Clinical isolates of carbapenem-resistant *K. pneumonae* (CRKP) were also selected to test the activity of 6D1 and kanamycin combination. The results showed that 6D1 did not reverse the antibacterial activity of kanamycin in CRKP containing the kanamycin resistance gene. However, we found that 6D1 enhanced the activity of kanamycin against a strain of CRKP that harbored the NDM-1 gene (Table 5, Table S1).

**Table 5.**
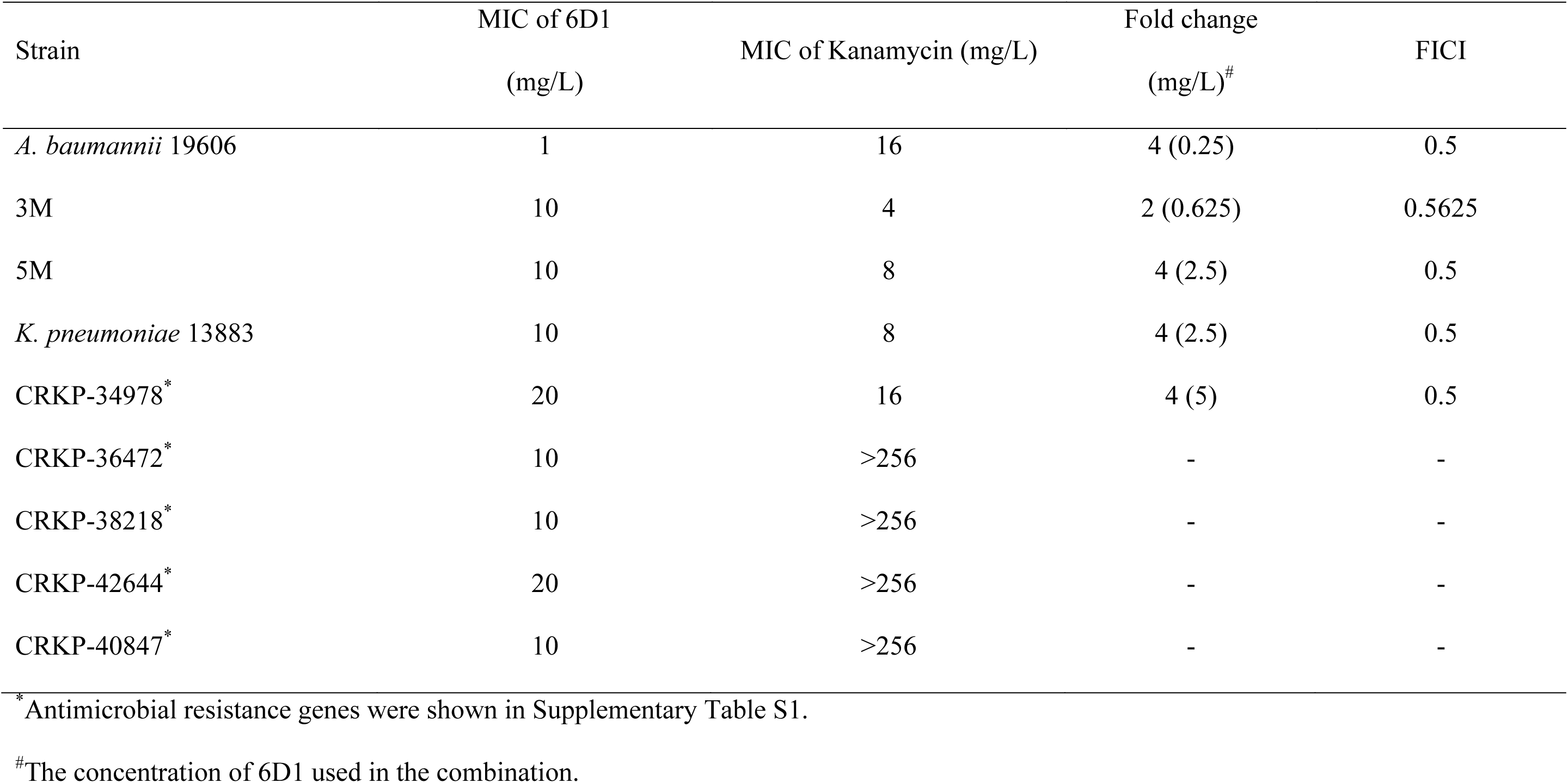
Potency of 6D1 in combination with kanamycin against different species

| Strain | MIC of 6D1<br>(mg/L) | MIC of Kanamycin (mg/L) | Fold change<br>(mg/L)<sup>#</sup> | FICI |
| --- | --- | --- | --- | --- |
| *A. baumannii* 19606 | 1 | 16 | 4 (0.25) | 0.5 |
| 3M | 10 | 4 | 2 (0.625) | 0.5625 |
| 5M | 10 | 8 | 4 (2.5) | 0.5 |
| *K. pneumoniae* 13883 | 10 | 8 | 4 (2.5) | 0.5 |
| CRKP-34978<sup>*</sup> | 20 | 16 | 4 (5) | 0.5 |
| CRKP-36472<sup>*</sup> | 10 | >256 | - | - |
| CRKP-38218<sup>*</sup> | 10 | >256 | - | - |
| CRKP-42644<sup>*</sup> | 20 | >256 | - | - |
| CRKP-40847<sup>*</sup> | 10 | >256 | - | - |

<sup>\*</sup>Antimicrobial resistance genes were shown in Supplementary Table S1.
<sup>#</sup>The concentration of 6D1 used in the combination.

### The interaction between 6D1 and OhrB

To characterize the binding of 6D1 and OhrB, we first obtained the purified His-tag-fused recombinant OhrB protein. The interaction between 6D1 and OhrB was confirmed by SPR analysis. It demonstrated the binding of 6D1 to OhrB, with an association rate constant of *k*^*a*^ 2.33 × 10^3^ M^−1^s^−1^, a dissociation rate constant *k*_*d*_ 2.28 × 10^−3^ s^−1^, and an equilibrium dissociation constant *K*_*D*_ 9.79 × 10^−6^ M (Figure 3).

**Figure 3.**
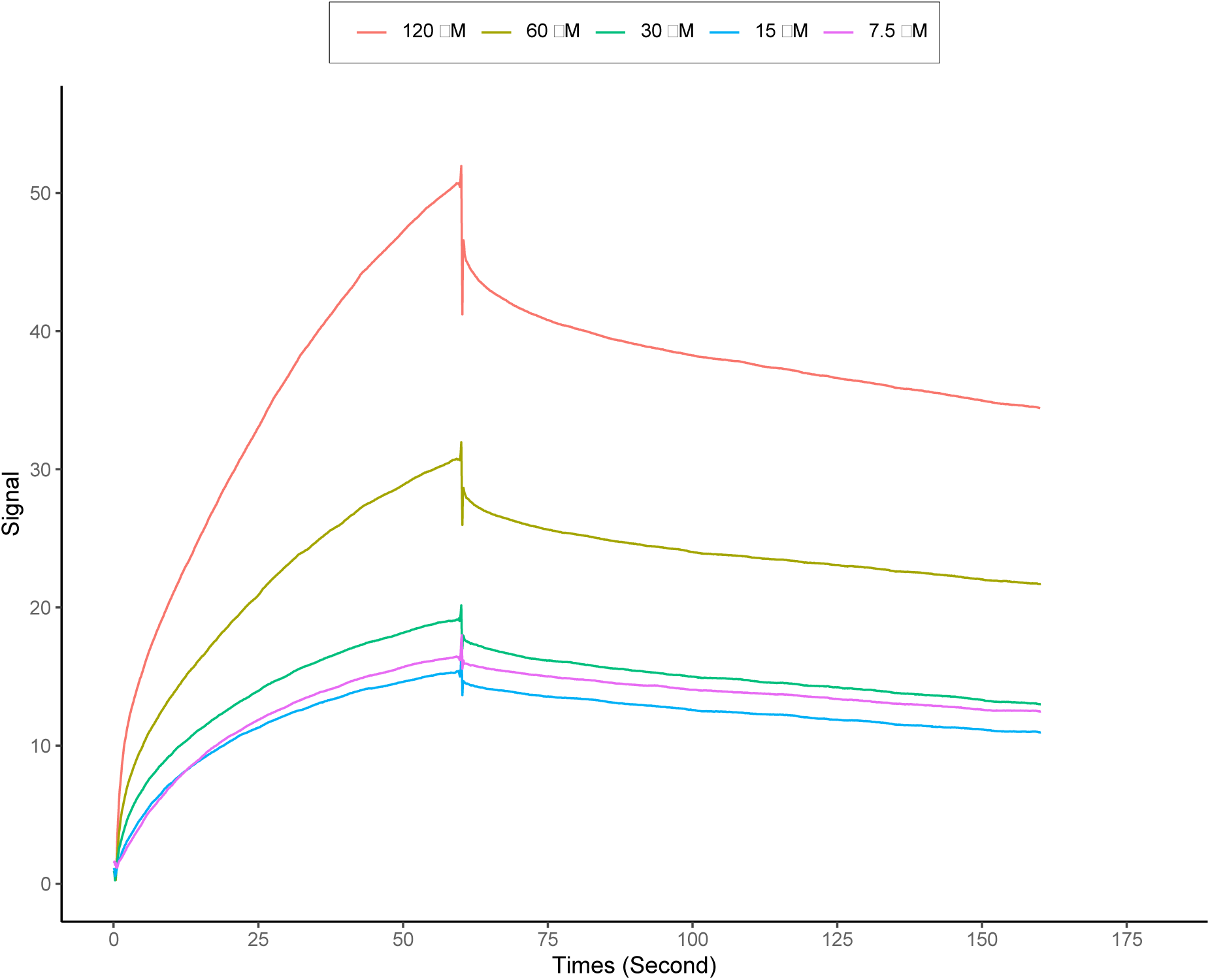
Surface plasmon resonance analysis of the interaction between 6D1 and OhrB. The *K*_*D*_ values were determined from the ratio between the kinetic rate constants (*k*_*a*_/*k*_*d*_).

## Discussion

Bacteria have evolved complex mechanisms to detoxify reactive oxygen species and thus strictly control hydroperoxide levels. A 9,540-bp plasmid pMAC carried by *A. baumannii* ATCC19606 that contains an OhrB coding region conferred bacterial resistant to organic peroxide-generating compounds CHP and *t*-BHP was reported in 2006.^17^ In this study, a whole-cell assay revealed a compound 6D1, which imparted inhibitory effects on *A. baumannii* ATCC19606 and had a similar structure to BIT and the antifungal ticlatone. The mutation site of the spontaneously resistant mutant suggested that the target of 6D1 was pMAC-derived OhrB. A previous study suggested that cellular thiol groups are major targets of BIT.^18^ Therefore, it provides a rationale that 6D1 acts on the thiol groups of OhrB. This is concordant with our result that 6D1 sensitizes *A. baumannii* ATCC19606 to CHP and *t*-BHP. Because most clinical isolates do not contain pMAC, this can explain why 6D1 is ineffective in clinical isolates.

In addition to developing antibiotics with new chemical structures and acting mechanisms, antibiotic adjuvants offer an alternative approach to combat resistance.^19^ In this study, 6D1 was found to impart an inhibitory effect on OhrB, and thus it is reasonable to use this as an adjuvant in combination with other antibiotics that induce bacteria to produce hydroperoxides. In addition, a previous study showed that all bactericidal antibiotics induce protective responses to reactive oxygen species.^20^ This suggests the potential of 6D1 as an adjuvant for bactericidal drugs. However, our data showed that 6D1 only enhances the activity of the aminoglycoside drug kanamycin but not others. This may be related to the reactivity order of Ohr to different peroxides, in which it mainly modulates the levels of fatty acid hydroperoxides and peroxynitrite.^16^ Because the effect of 6D1 is achieved by inhibiting OhrB, it is not surprising that 6D1 was not effective on drug-resistant strains that harbored kanamycin resistance genes such as the 16s rRNA methylase enzyme *rmtB*. Notably, the combination of 6D1 and kanamycin was effective on the CRKP strain carrying NDM-1. A recent study has shown that in Northeast China, the aminoglycoside resistance gene *rmtB* was detected in 96.61% of KPC-2-producing CRKP and in 21.74% of NDM-1-producing CRKP, indicating the potential combinative application of kanamycin and a peroxidase inhibitor such as 6D1 in about 80% of NDM-1-producing CRKP.^21^

The clinical use of kanamycin has been limited by its well-known toxicity and side effects such as ototoxicity. Our study revealed the feasibility of enhancing the activity of kanamycin by inhibiting the detoxification ability of bacteria to peroxides, thereby providing a new target and strategy for the development of kanamycin enhancers in the near future.

## Transparency declarations

None to declare.

---

**Table S1.**
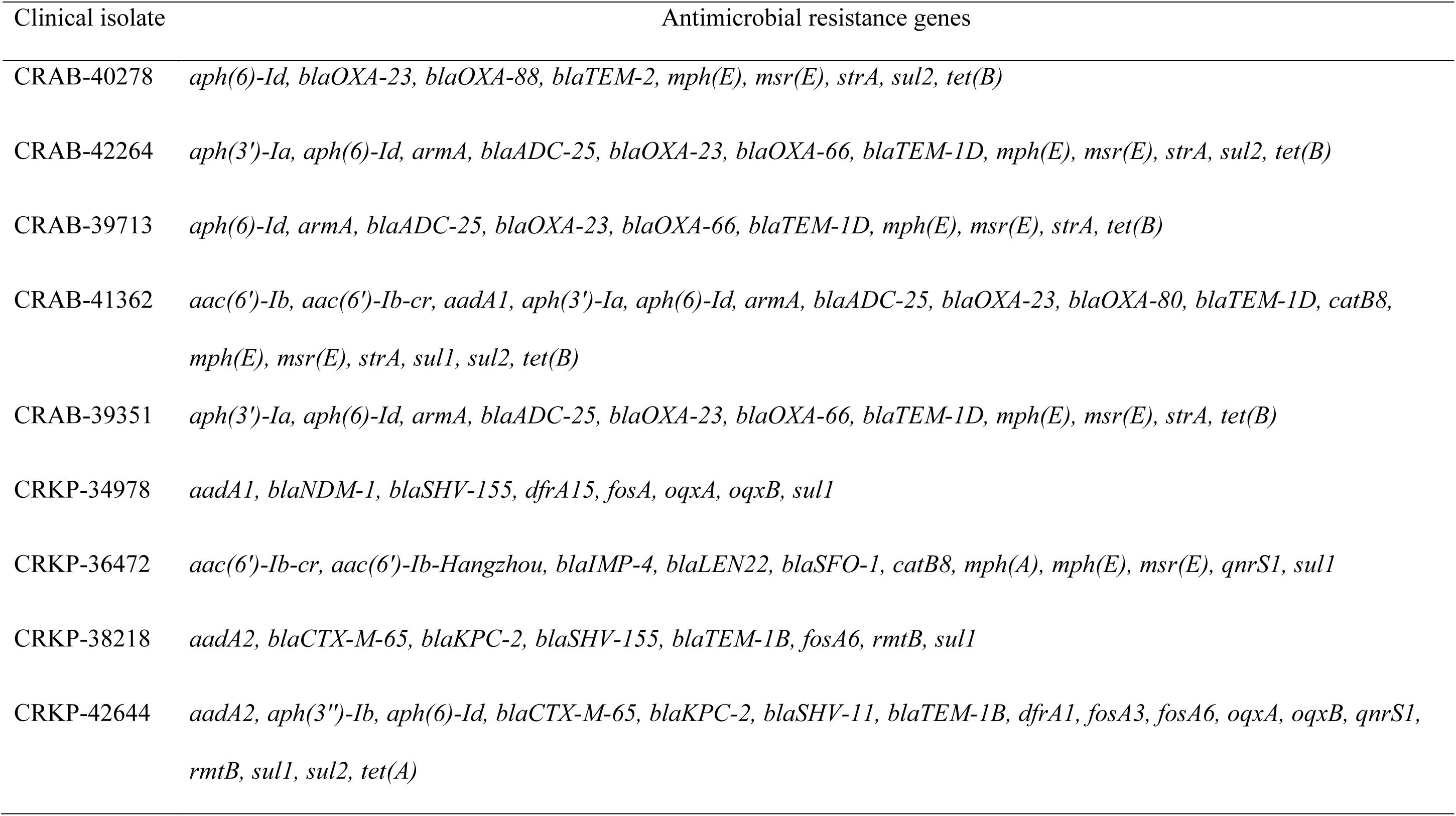
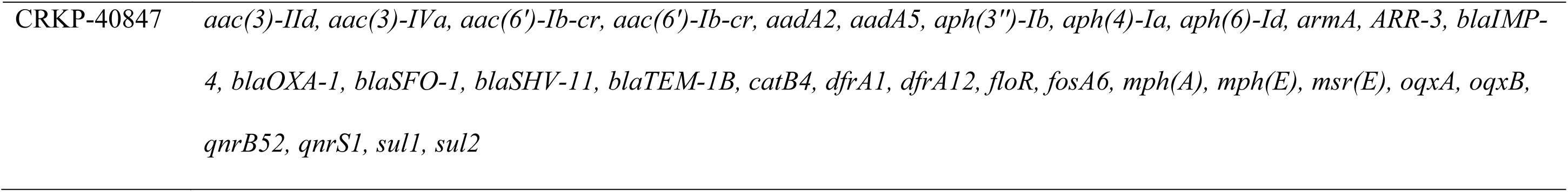
Antimicrobial resistance genes in clinical isolates

